# Redundant YAP and TAZ functions are essential for skeletal muscle regeneration

**DOI:** 10.64898/2026.09.04.749462

**Authors:** Jason S. Silver, Alicia A. Cutler, Tze-Ling Chang, Kristi S. Anseth, Bradley B. Olwin

## Abstract

Muscle stem cells orchestrate skeletal muscle regeneration through complex fate decisions. The transcriptional co-activators Yes-associated protein 1 (YAP) and WW domain-containing transcription regulator 1 (TAZ) contribute to multiple stages of myogenesis, yet their individual contributions to regeneration remain unclear due to substantial functional overlap. We genetically titrated YAP and TAZ expression in MuSCs with double knockout and single allele mutants by crossing Pax7^CreERT^ mice with TAZ^flox/flox^;YAP^flox/flox^ mice. Conditional deletion of both YAP and TAZ in muscle stem cells severely disrupted muscle regeneration with dramatically increased fibrosis and impaired myofiber formation following injury. In contrast, a single allele of either YAP or TAZ was sufficient to rescue injured muscle weight and myofiber cross-sectional area. Similarly, the reduced proliferation of double knockout muscle stem cells on isolated myofibers was restored by a single allele of either YAP or TAZ. In addition, disrupted actin cytoskeleton organization and reduced focal adhesion formation drove double knockout muscle stem cell migration defects, negatively impacting muscle stem cell congregation prior to fusion. Thus, YAP and TAZ function redundantly as critical transcriptional co-activators to regulate progenitor proliferation and migration during muscle regeneration. By challenging myogenesis with double knockout of both YAP and TAZ, we unmasked regenerative requirements previously undetected in single-gene loss models, highlighting genetic redundancy as a key principle buffering regenerative robustness.

## Introduction

Skeletal muscle is composed of post-mitotic, multinucleated myofibers that depend on resident muscle stem cells (MuSCs) to maintain and repair muscle tissue. Upon damage, quiescent MuSCs activate, executing a coordinated sequence of cellular events with cells migrating, proliferating, committing to differentiate, and ultimately fusing to repair myofibers^1^. Coordinated signaling from extrinsic niche cues and intrinsic pathways orchestrates MuSC cell fate decisions during muscle repair^2,3^. Although many exogenous cues and individual signaling pathways controlling discrete aspects of MuSC behavior have been characterized, how molecular regulators collectively drive cell fate decisions to regenerate muscle remains incompletely defined.

Among candidate regulators of MuSC fate are the transcriptional co-activators Yes-associated protein 1 (YAP) and WW domain-containing transcriptional regulator 1 (WWTR1 or TAZ). YAP and TAZ regulate cell cycle progression, cytoskeletal organization, motility, and cell growth, features essential for coordinating stem cell behavior during regeneration^4^. In skeletal muscle, both proteins are involved in activating MuSCs^5^, expanding MuSCs^6–8^, and expanding C2C12 myoblasts^8^, as well as differentiating MuSCs^7^ and C2C12 myoblasts^9,10^. In MuSCs, both proteins are upregulated upon activation^5^, maintained during cell expansion^7,9^, and deactivated as cells exit the cell cycle^9,10^, consistent with roles in promoting activated and proliferating MuSCs rather than regulating commitment of MuSCs to a terminal post-mitotic fate. YAP and TAZ are mechanotransducers, translating extracellular matrix mechanics into cellular responses by regulating their subcellular localization via LATS1/2 kinases-mediated phosphorylation through the Hippo signaling pathway^4,7^. Ablating both YAP and TAZ in MuSCs after injury prevented MuSCs from expanding in response to pathologically stiff microenvironments^8^. Although YAP and TAZ regulate aspects of myogenesis, their individual functions in directing MuSC behavior in regenerating muscle remain unknown.

YAP and TAZ are paralogs that share ∼40% amino acid sequence homology, similar protein-protein interaction motifs, and largely convergent regulatory mechanisms^11,12^. Their functional overlap has complicated efforts to define the individual contributions of each transcriptional co-activator in regenerating muscle. Deletion of either YAP or TAZ in MuSCs produced only modest effects on the ability of skeletal muscle to regenerate after injury^7,13,14^. Embryonic deletion of both YAP and TAZ in myofibers disrupted developmental myogenesis, leading to diaphragm insufficiency and postnatal arrest^15^. In contrast, mice that retained either YAP or TAZ in myofibers survived to adulthood with body weights comparable to controls, but reduced grip strength, indicating partial functional redundancy of the two proteins^15^. While YAP and TAZ embryonic deletion caused perinatal lethality, YAP induced myofiber hypertrophy following denervation^15,16^. The extent that YAP and TAZ act redundantly in MuSCs, and whether this redundancy masks essential regenerative functions, remain unresolved.

We directly tested whether YAP and TAZ individually are dispensable for muscle repair by genetically titrating YAP and TAZ dosage in MuSCs as muscles regenerate. Using conditional single-allele and double-allele deletion, we determined whether their combined activity is required for effective skeletal muscle regeneration after injury. Our findings reveal that YAP and TAZ function redundantly yet indispensably in MuSCs where combined loss profoundly disrupts regenerative capacity through defects in proliferation and migration. These results establish that YAP and TAZ functions required for skeletal muscle regeneration are masked by genetic compensation.

## Results

### YAP and TAZ redundantly support skeletal muscle regeneration

YAP and TAZ regulate MuSCs with some overlapping functions, but the individual contributions of each transcriptional co-activator as well as their combined roles have not yet been tested. To test the potential overlapping and individual effects, we generated an allelic series of MuSC-specific, conditional YAP and TAZ knockout mice by breeding Pax7^CreERT^ mice^17^ with TAZ^flox/flox^;YAP^flox/flox^ mice^18^. Cre-mediated recombination induced by administering tamoxifen excises YAP and TAZ to generate double knockout (dKO: TAZ^-/-^; YAP^-/-^; Cre^+^) MuSCs as well as single allele (TAZ SA: TAZ^+/-^; YAP^-/-^; Cre^+^ and YAP SA: TAZ^-/-^; YAP^+/-^; Cre^+^) MuSCs (**Fig. 1A**). We assessed the effects of allelic YAP and TAZ combinations on the capacity of MuSCs to regenerate muscle by injuring tibialis anterior (TA) muscles with barium chloride (BaCl_2_) and assaying at 28 days post-injury (DPI), when myofibers are morphometrically and functionally recovered (**Fig. 1B**). TA muscles in dKO mice failed to regenerate, as fewer regenerated myofibers containing centrally located nuclei were present accompanied by large fibrotic lesions compared to control mouse TA muscles lacking Cre recombinase (Cre Null: Cre^-/-^) (**Fig. 1C**). The mass of the injured TA muscles relative to the contralateral uninjured muscles was reduced by approximately 50% in dKO mice, consistent with a failure to regenerate (**Fig. 1D**). The minimum Feret diameters of regenerated myofibers in dKO TA muscles were smaller compared to wild type TA muscles (**Fig. 1E**). The few remaining uninjured myofibers in dKO TA muscles were hypertrophic, suggesting compensatory growth in response to failed repair (**Fig. 1F**). A single YAP or TAZ allele in MuSCs partially rescued muscle repair, as the normalized injured TA muscle mass (ratio of injured TA muscle mass/contralateral TA muscle mass) in either YAP SA or TAZ SA mice was comparable to the normalized injured TA muscle mass in Cre Null control mice (**Fig. 1C,D**). The regenerated TA muscles were comprised of abundant centrally nucleated myofibers (**Fig. 1C**). Although the normalized TA muscle mass in single-allele knockout mice was comparable to that of Cre Null control mice, the minimum Feret diameters of regenerated myofibers in TA muscles from YAP SA (mean 37.8 ± 7.7 μm) and TAZ SA (mean 37.9 ± 9.7 μm) mice were larger than those from dKO mice (mean 30.7 ± 10.7 μm), but smaller than those from Cre Null control mice (median 46.5 ± 10.4 μm) (**Fig. 1E**). Thus, mice with a single allele of either YAP or TAZ are capable of a regenerative response but display deficits compared to Cre Null control mice.

**Figure 1:**
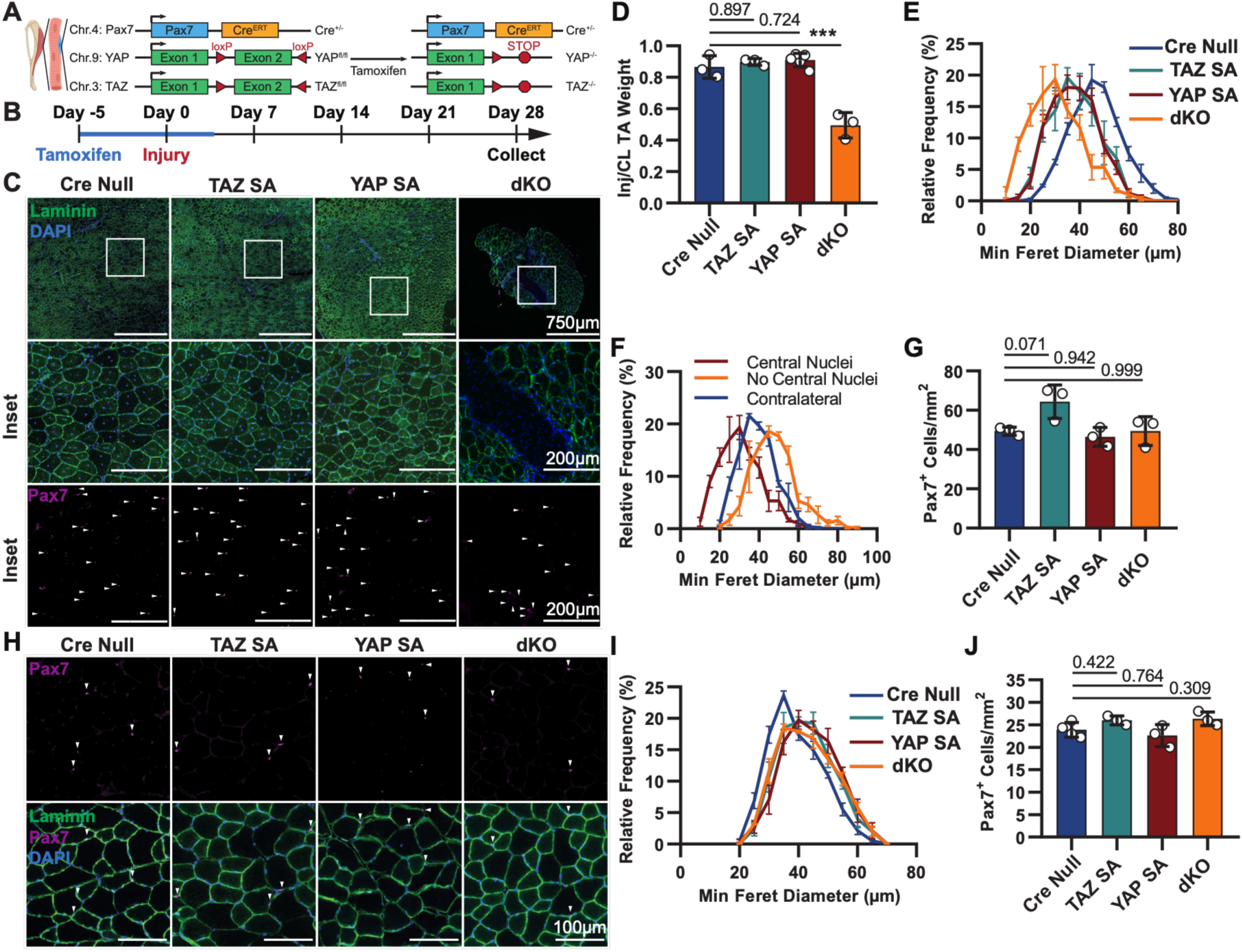
YAP and TAZ are essential for muscle regeneration. A) Schematic of MuSC-specific conditional YAP and TAZ knockout (YAP^fl/fl^; TAZ^fl/fl^; Pax7^CreERT^) mice. B) Experimental timeline depicting tamoxifen administered for 5 days to induce recombination in MuSCs, followed by BaCl_2_-induced injury of the TA muscles and tissue collection for histological analysis at 28 days post-injury (DPI). C) Representative histological sections are assayed for immunoreactivity to laminin and Pax7 with nuclei labeled with DAPI. White arrowheads identify Pax7^+^ MuSCs and insets are higher-magnification views of regenerated regions. D) The dissected injured (Inj) TA muscle weight was normalized to the uninjured contralateral (CL) muscle. E) The distribution of minimum Feret diameter of myofibers identified by laminin immunoreactivity. Graph displays mean ± standard error. F) The distribution of minimum Feret diameters of myofibers in injured muscles from dKO mice, classified by the presence or absence of centrally located nuclei, compared to myofibers in uninjured, contralateral TA muscles. G) The density of Pax7^+^ MuSCs normalized to muscle area is quantified in 28 DPI TA muscle sections. H) Representative histological sections of uninjured contralateral TA muscles assayed for immunoreactivity to Pax7 and laminin. The minimum Feret diameter (I) and density of Pax7^+^ MuSCs (J) of contralateral muscles are plotted. Unless noted elsewhere, at least 3 biological replicates were analyzed per condition with >50 MuSCs quantified per mouse. Error bars represent the SD and ***p<0.001 by one-way ANOVA.

Ablating YAP and TAZ in MuSCs compromises the regenerative capability of the stem cells. To better understand the cellular processes involved, we examined how the loss of YAP, TAZ, or both factors impacts the MuSC population. At 28 days post-injury, the density of Pax7^+^ MuSCs in TA muscles from dKO mice was not significantly different from that observed in Cre Null or YAP SA mice (**Fig. 1G**). However, Pax7^+^ MuSCs did not occupy the MuSC niche in dKO mice and were instead located within the TA muscle interstitial space (**Fig. 1C**). Unexpectedly, Pax7^+^ MuSC density in TA muscles from TAZ SA mice was greater than in all other genotypes (**Fig. 1G**). Given the effects observed on MuSCs and their ability to repair muscle by altering YAP or TAZ dosage, we asked if loss of YAP, TAZ, or both alleles affects TA muscles in the absence of injury. Myofiber diameter and MuSC density in uninjured TA muscles were unaffected by ablating either YAP, TAZ, or both alleles (**Fig. 1H-J**). Thus, YAP and TAZ appear dispensable for short-term muscle homeostasis, and a single allele of either YAP or TAZ is sufficient to regenerate TA muscles following an induced injury, but loss of both YAP and TAZ alleles severely compromises the ability of MuSCs to repair muscle.

### Loss of YAP and TAZ impairs MuSC proliferation

A single allele of either YAP or TAZ partially rescues muscle regeneration by MuSCs, whereas double knockout of both alleles in MuSCs prohibits regeneration. To determine whether deleting YAP and TAZ is promoting MuSC apoptosis, we cultured MuSCs from dKO mice, induced recombination *ex vivo,* and evaluated apoptotic DNA cleavage using a TUNEL assay (**Fig. 2A**). Fragmented DNA was not observed in dKO or control cultures but was prevalent when MuSCs were treated with DNase I, confirming assay sensitivity (**Fig. 2B, C**). Loss of YAP and TAZ does not appear to promote MuSC apoptosis and instead is likely directly affecting the capability of MuSCs to repair muscle tissue.

**Figure 2:**
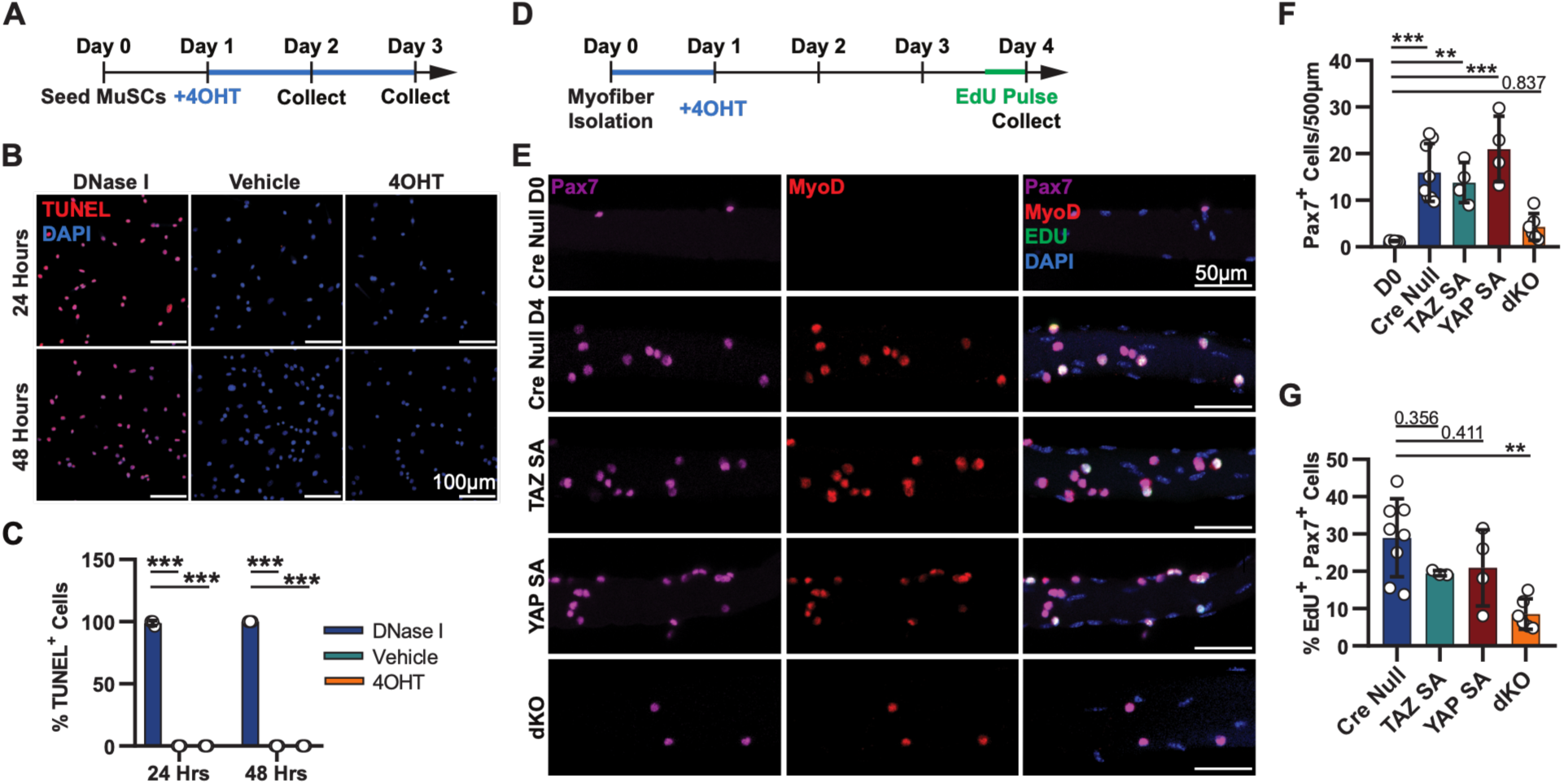
YAP and TAZ knockout reduces MuSC proliferation sparing MyoD activation. A) Experimental schematic showing *ex vivo* MuSC culture, induction of recombination with 4OHT one day after seeding, and collected at 24 and 48 hours. B) Representative images of MuSCs assayed for DNA fragmentation using the TUNEL assay. Nuclei are detected by DAPI. C) Quantification of TUNEL^+^ cells is plotted. D) Schematic of myofiber-associated MuSC proliferation assay. EDL myofibers were isolated, treated with 4OHT to induce recombination, and cultured for 4 days before receiving a 2-hour EdU pulse prior to harvest. E) Representative images of MuSCs on myofibers assayed for EdU incorporation and for immunoreactivity to Pax7 and MyoD. Quantification of Pax7^+^ MuSC number (F) and percentage of EdU^+^ MuSCs (G) after 4 days in culture. n = 3 biological replicates. Error bars represent the SD and **p<0.01, ***p<0.001 by one-way ANOVA.

To repair muscle, MuSCs exit quiescence and transition through several cell fate changes, eventually fusing with each other or fusing with existing myofibers. To assess the effects of YAP and TAZ loss on these cell fate transitions in dKO mice, we isolated single myofibers with associated resident MuSCs from extensor digitorum longus (EDL) muscles. The cultures were treated with 4-hydroxytamoxifen (4OHT) immediately upon isolation when MuSCs were already activated^19,20^, and at four days in culture, proliferating MuSCs were treated with 5-ethynyl-2’- deoxyuridine (EdU) to identify MuSCs in S-phase, with cultures fixed and proliferating cells quantified (**Fig. 2D,E**). Wild type MuSCs expanded robustly relative to initial MuSC numbers on freshly isolated intact myofibers (**Fig. 2F**). In dKO EDL muscles, Pax7+/EdU+ MuSC numbers were reduced three-fold, while Pax7+/EdU+ MuSC numbers on intact myofibers expressing a single allele of either YAP or TAZ were indistinguishable from those in wild type (Cre null) mice (**Fig. 2G**).

### YAP and TAZ are dispensable for myogenic commitment of MuSCs but regulate density-dependent MuSC fusion

MuSCs activate when isolated, rapidly inducing MyoD protein that commits MuSCs to the myogenic lineage^21–24^. MyoD expressing myoblasts can either self-renew and re-acquire quiescence to re-establish the MuSC pool, proliferate as myoblasts, or induce Myogenin to terminally differentiatiate^25–28^. To assess whether loss of YAP and TAZ affects MuSC fate transitions, we assayed immunoreactivity of MyoD and Myogenin in MuSCs on intact myofibers from wild type and dKO mice. Single myofibers and their associated MuSCs were isolated from tamoxifen-injected mice and cultured for 6 days, with myofibers fixed daily and assayed for MyoD and Myogenin immunoreactivity (**Fig. 3A**). Syndecan-4 immunoreactivity was used to identify MuSCs as Pax7 protein is lost when the cells terminally differentiate. The fraction of Syndecan-4^+^/MyoD^+^ MuSCs was comparable in wild type and dKO mice as well as in mice expressing a single allele of either YAP or TAZ in MuSCs (**Fig. 3B,C**). Commitment to terminal differentiation, assessed by Myogenin immunoreactivity, emerged after 2 days of culture, with nearly all MuSCs of any genotype positive for Myogenin immunoreactivity at 6 days of culture (**Fig. 3B,C**). Ablating both YAP and TAZ elicited no significant changes in the timing of appearance for myogenic transcription factor proteins in MuSCs cultured on intact myofibers, despite the lower total number of dKO MuSCs relative to wild type MuSCs in Cre Null mice (see **Fig. 2F**).

**Figure 3:**
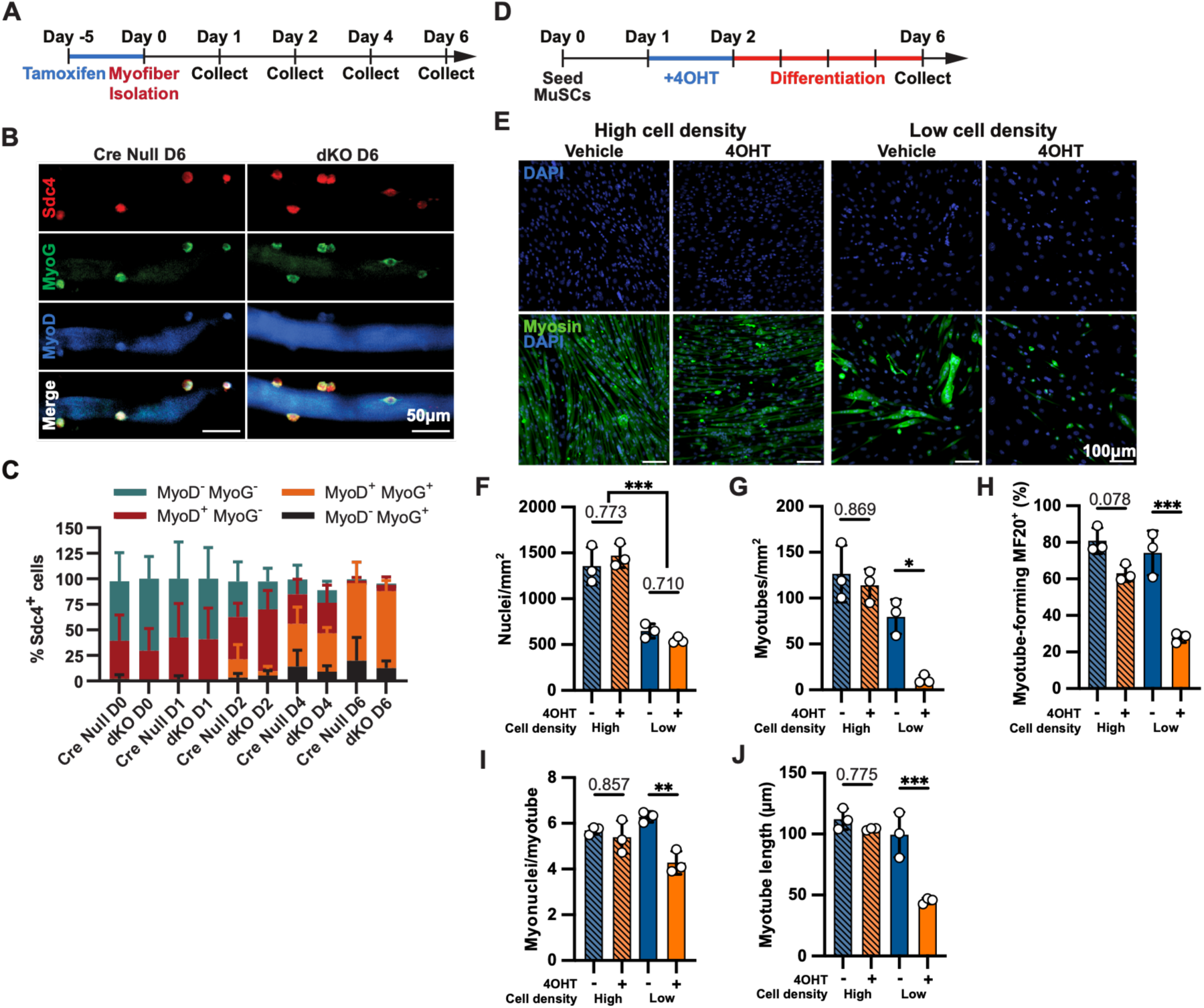
MuSC differentiation is impacted by YAP and TAZ in a density-dependent manner. A) Experimental schematic depicting *in vivo* tamoxifen treatment prior to myofiber harvest. B) Isolated myofibers cultured in suspension were assayed for MuSCs (Sdc4+) and quantified (C) for immunoreactivity to MyoD and MyoG over 6 days post-isolation. D) Schematic of *in vitro* density-dependent differentiation assay where MuSCs from dKO mice were seeded at high and low density and treated with either 4OHT or vehicle for 24 hours before inducing differentiation with serum starvation. E) Representative images of differentiated myotubes were detected by immunoreactivity to MF20. The densities of nuclei (F) and myotubes (MF20+ syncytia with more than 3 nuclei) (G) were quantified. The percentage of MF20+ myocytes within multinucleated myotubes was calculated (H). The average myonuclear number (I) and myotube length (J) were quantified. At least 3 biological replicates were analyzed per condition. Error bars represent the SD and *p<0.05, **p<0.01, ***p<0.001 by one-way ANOVA.

The failure of dKO MuSCs to expand is not associated with altered timing of myogenic proteins regulating MuSC fate, suggesting that the failure of MuSCs to expand occurs from yet undiscovered cellular deficits. Since YAP and TAZ transduce mechanical cues into chemical responses and MuSCs on intact myofibers are initially sparse but expand into dense colonies, we asked if cell density in culture affects dKO MuSC behavior. To assess if cell density differentially affects dKO MuSCs compared to wild type MuSCs, MuSCs were isolated from dKO mice, seeded at either high or low density and treated with 4OHT or vehicle to generate dKO or control MuSCs *ex vivo*. The cultures were then induced to differentiate by reducing serum (**Fig. 3D,E**), cultured for 4 days, and assessed for nuclear density (**Fig. 3F**), myosin-immunoreactive multinucleated myotubes (**Fig. 3G,H**), frequency of myonuclei (**Fig. 3I**), and myotube length (**Fig. 3J**). Low-density cultures induced to differentiate possessed similar nuclear densities (**Fig. 3F**), but dKO cultures had dramatically fewer myotubes and differentiated cells (**Fig. 3G,H**), fewer myonuclei per myotube (**Fig. 3I**), and decreased myotube length (**Fig. 3J**) compared to wild type cultures. The capacity of dKO MuSCs to differentiate is severely affected by cell density where dKO differentiate poorly at low cell density but differentiate similarly to wild type MuSCs when cultured at high cell densities.

### YAP and TAZ regulate MuSC migration through cytoskeletal and focal adhesion organization

Deleting YAP and TAZ from fibroblasts and endothelial cells affects their ability to migrate^29,30^, suggesting that dKO MuSCs at low density may have impaired cell-cell interactions, limiting their ability to differentiate and fuse. The motility of MuSCs isolated from dKO mice was tracked via live-cell imaging over 24 hours in culture (**Fig. 4A**) where migratory speed and cell trajectories were dramatically altered in dKO MuSCs compared to wild type MuSCs (**Fig. 4B**). The mean speed (**Fig. 4C**) and trajectory straightness (**Fig. 4D**) were significantly lower in dKO MuSCs compared to wild type MuSCs.

**Figure 4:**
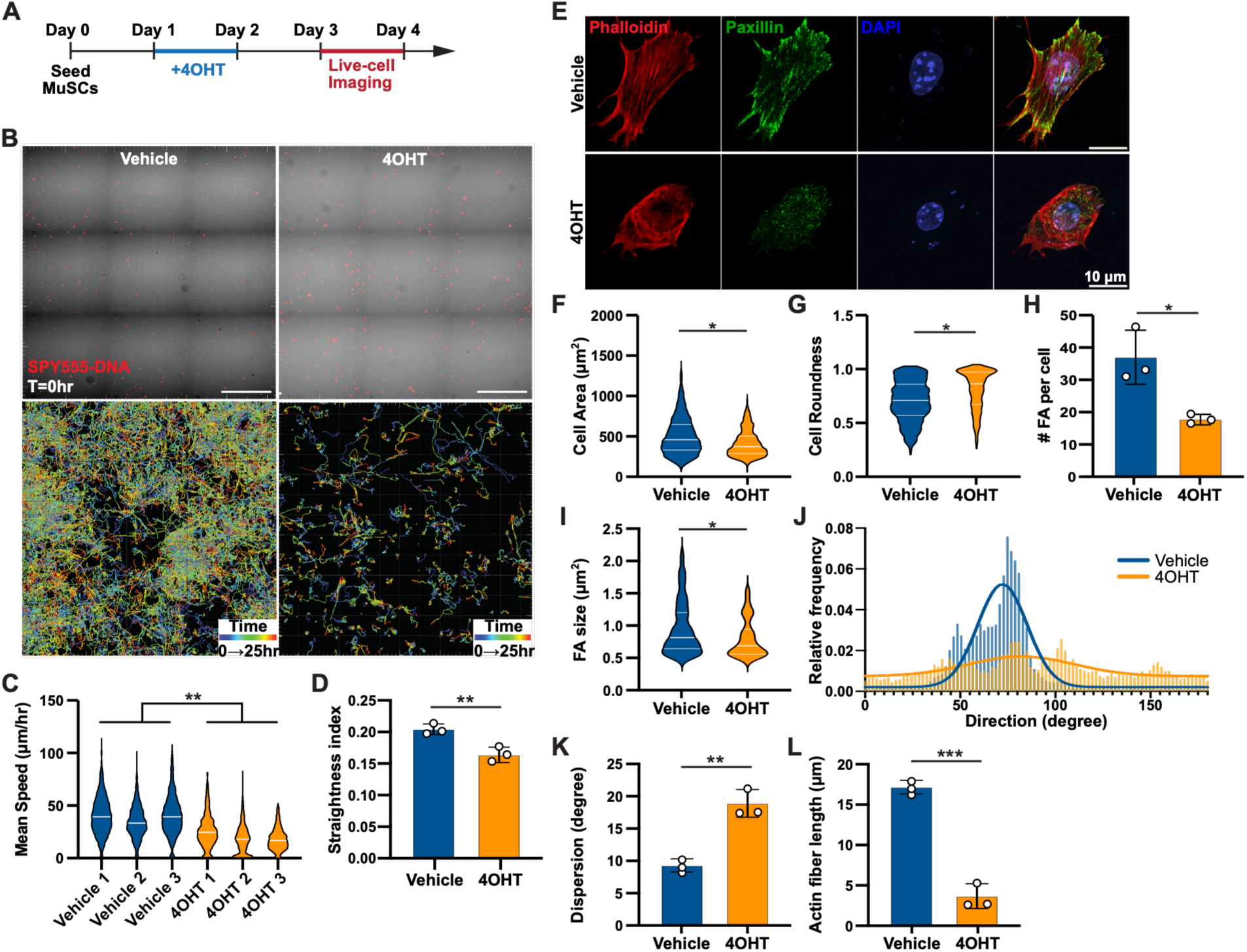
Loss of YAP and TAZ impairs MuSC migration via disruption of cytoskeletal and focal adhesion organization. A) Schematic of live-cell migration assay where MuSCs were fluorescently labeled with SPY555-DNA and positionally tracked over 24 hours with image acquisition every 15 minutes following recombination. B) Representative migration tracks of individual MuSCs over a 24-hour imaging period with quantification of migration speed (C). D) Straightness index is calculated as the ratio of the net displacement to the total path length. E) Representative images of MuSCs cultured *in vitro*, treated with 4OHT or vehicle 24 hours post-seeding, fixed 2 days post-4OHT removal, and immunoassayed for F-actin via phalloidin and focal adhesions via paxillin. Quantification of projected cell area (F) and cell roundness (G) were plotted. Focal adhesion number per cell (H) and size (I) were quantified. Histogram of actin fiber alignment (J) and average dispersion (K) were plotted along with average fiber length (L). At least 3 biological replicates were analyzed per condition. Error bars represent the SD and *p<0.05, **p<0.01, ***p<0.001 by one-way ANOVA.

To determine whether the loss of YAP and TAZ affects the cell morphology, actin cytoskeleton, and focal adhesions^31,32^, we assessed phalloidin staining and immunoreactivity for Paxillin in wild type and dKO MuSCs. Long cellular protrusions and focal adhesions apparent in wild type MuSCs were generally absent in dKO MuSCs (**Fig. 4E**). When quantified, dKO MuSC projected areas and roundness were significantly different relative to wild type MuSCs, where dKO MuSCs were smaller with higher roundness indicating reduced cell spreading (**Fig. 4F,G**). The number and size of immunoreactive paxillin focal adhesions were significantly reduced in dKO MuSCs compared to wild type MuSCs, suggesting a failure of focal adhesions to mature (**Fig. 4H,I**). Actin fibers were significantly shorter and less aligned in dKO MuSCs compared to wild type MuSCs (**Fig. 4J-L**), reflecting an altered cytoskeleton and suggesting that YAP and TAZ are essential for regulating MuSC motility.

## Discussion

Skeletal muscle is continuously maintained by MuSCs that must exit quiescence, migrate, and either expand before fusing into existing myofibers or fuse without an intervening cell cycle^15,16^. Since MuSCs are sparse in muscle tissue, comprising only a small percent of the total nuclei, MuSCs likely migrate to maintain and repair muscle tissue. We found that YAP and TAZ are essential for MuSCs to appropriately repair and maintain muscle, likely transducing mechanical cues into signaling pathways that regulate MuSC movement required for differentiation and fusion. Although incompletely redundant, a single allele of either YAP or TAZ rescues migratory and fusion deficits in MuSCs that occurr when both alleles are ablated. Either YAP or TAZ allele is capable of sustaining the transcriptional output required to regenerate muscle, explaining why prior single-gene knockout studies reported only modest phenotypes^7,13–15^. Functional redundancy among paralogous transcriptional regulators is a potential mechanism to buffer tissue repair against genetic or environmental perturbations^33^. Similar compensatory mechanisms within myogenic regulatory networks occur in *C. elegan* where HLH-1, UNC-120, and HND-1 redundantly specify body wall muscle differentiation; loss of any one or two of these factors has a minimal effect on morphogenesis but deleting all three factors prevents body wall myogenesis^34^.

A single YAP or TAZ allele is required to regenerate muscle, yet YAP and TAZ exert distinct biological roles in development and tissue homeostasis^12^. Germline deletion of YAP results in developmental arrest and embryonic lethality, whereas TAZ knockout mice survive to adulthood but develop renal and pulmonary pathologies^35–37^. Expression of either YAP or TAZ in myofibers is sufficient to rescue perinatal lethality; however, the loss of TAZ has a more pronounced impact on postnatal growth and grip strength than the loss of YAP^15^. In contrast to our data where we observed comparable effects with single alleles of either YAP or TAZ, specifically deleting YAP in MuSCs yielded mild regenerative defects, whereas deleting TAZ produced no observable impairment^7^. Chromatin immunoprecipitation sequencing of TEAD, the DNA-binding partner of YAP and TAZ, in immortalized myoblast progenitors indicates that TAZ more strongly drives transcriptional programs associated with cell cycle progression than YAP^15,38^. Thus, YAP and TAZ may differentially regulate MuSCs, but defining the context-specific transcriptional and epigenetic mechanisms that YAP and TAZ uniquely regulate will require additional experiments.

YAP regulates muscle hypertrophy^16^, yet uninjured myofibers in our dKO mice appear to hypertrophy and thus, we conclude that neither YAP or TAZ is required for hypertrophy; alternatively, residual YAP and TAZ protein present in myonuclei may be sufficient to sustain hypertrophy in short-term experiments. We then asked whether YAP or TAZ is necessary to maintain muscle in young mice and deleted YAP and TAZ in MuSCs assaying muscle tissue 30 days post-ablation. Uninjured muscles lacking YAP and TAZ in MuSCs were indistinguishable from wild type muscle indicating that loss of YAP and TAZ do not affect maintenance of muscle over 30 days but may be required for long-term muscle maintenance.

The failure to regenerate muscle is attributable, in part, to proliferative and migratory impairments in dKO MuSCs. A single allele of either YAP or TAZ largely restored MuSC-mediated muscle tissue repair, presumably by enabling MuSCs to migrate, differentiate, and fuse. It remains unclear whether the primary YAP and TAZ response in MuSCs arises from cytokine signaling via the Hippo pathway^39^, altered mechanotransduction signaling^8,40^, or both. MuSCs lacking both YAP and TAZ failed to migrate appropriately, exhibiting reduced motility and loss of directional persistence, accompanied by altered cell morphology, a disorganized actin cytoskeleton, and diminished focal adhesions. These phenotypes are consistent with altered MuSC mechanotransduction (e.g., cells perceiving a mechanically soft environment), despite being cultured on supraphysiologically stiff plastic^41^. Thus, YAP and TAZ sustain cytoskeletal tension and focal adhesion dynamics via matrix interactions that may involve increased expression of *TRIO*, a guanine nucleotide exchange factor activating Rac1 while suppressing RhoA GTPases to promote cell motility^42^. Ablating YAP and TAZ may uncouple environmental mechanics from intracellular organization, impairing MuSC motility and reducing MuSC regenerative capability. Delineating the mechanisms by which YAP and TAZ integrate mechanical and biochemical cues to permit attachment, migration, and fusion as muscle regenerates may explain in part why regeneration is compromised in pathologically stiffened muscle and help identify new therapeutic targets to restore MuSC function.

## Materials and Methods

### Animals

Mice were bred and housed according to National Institutes of Health guidelines for the ethical treatment of animals in a pathogen-free animal facility at the University of Colorado, Boulder. The University of Colorado Institutional Animal Care and Use Committee approved all animal protocols and ensured procedures complied with all ethical regulations. To generate a conditional knockout of YAP and TAZ, Pax7^CreERT^ mice^17^ were crossed into the YAP1^tm1Hmc^;WWTR1^tm1Hmc^ mice (Jackson Laboratory, Stock No: 030532)^18^. Genotyping was performed on tissue samples collected at weaning and sent to Transnetyx for automated genotyping. Control mice were age and sex matched.

### Animal Procedures

For barium chloride injuries, mice 3-6 months old were anesthetized using isoflurane, BaCl_2_ solution (50 μL, 1.2% in saline) was then injected into the tibialis anterior muscle^43^. For genetic knockout, tamoxifen (Sigma) was resuspended in corn oil (Sigma) and administered by daily intraperitoneal injections (0.075 mg/gram) for 5 days. Mice were fed a diet containing tamoxifen (250 mg/kg, Envigo) for 3 days post-injury.

### Histology and Immunohistochemistry

Muscle histology was conducted on the tibialis anterior (TA) muscles. Mice were euthanized according to NIH guidelines and the muscles isolated. TA muscles were fixed for 2 hours in paraformaldehyde (4%) on ice and then dehydrated in sucrose (30%) overnight at 4°C. Tissues were mounted in O.C.T compound (Tissue-Tek) and frozen on dry ice. Cryo-sectioning was performed (Leica) to generate 10 μm sections. Muscle sections were fixed for 7 minutes in paraformaldehyde (4%) and then washed 3x in phosphate-buffered saline (PBS) for 10 minutes. Heat-induced epitope retrieval was performed for Pax7 immunodetection by transferring samples in citrate buffer (pH 6.0) and subjecting to high-pressure cooking (model CPC-600, Cuisinart) for 6 minutes. Tissue sections were cooled, washed 3x in PBS for 10 minutes, and permeabilized in blocking buffer (0.25% Triton X-100, 3% bovine serum albumin, Sigma) for 1 hour. Sections were then incubated with primary antibody cocktails in blocking buffer for 1 hour at room temperature. Prior to secondary staining, samples were washed 3x in PBS plus Triton X-100 (PBST, 0.1%, Sigma) for 10 minutes each. Samples were incubated with secondary antibody cocktails in blocking buffer for 1 hour at room temperature and washed 3x in PBST for 10 minutes. Tissue samples were mounted in Mowiol supplemented with DABCO (Sigma) and stored at room temperature until imaging. The following antibodies were used in these studies: anti-Pax7 (Developmental Studies Hybridoma Bank at 1:1000), anti-laminin (Sigma L9393, 1:200), and Alexa Fluor -488 and -647 conjugated secondary antibodies (Invitrogen at 1:500). DAPI (1μg/mL, Sigma) was used to detect nuclei.

### Myofiber isolation and culture

Mice were euthanized according to IACUC guidelines and the extensor digitorum longus (EDL) muscles were dissected. EDL muscles were enzymatically digested in collagenase (400 U/mL Worthington) in Ham’s F-12C (Gibco) for 90 min at 37°C on a rotating mixer and then quenched with 15% horse serum (Gibco) in Ham’s F-12C (Gibco) media. Using a flame-polished glass pasteur pipette, individual myofibers were manipulated and isolated from the digest. FGF (0.5 nM) was supplemented in media and myofibers were cultured in 5% O_2_ and 5% CO_2_ at 37°C for the indicated durations prior to fixation. For *ex vivo* genetic knockout, 4-hydroxytamoxifen (4OHT) (500 nM in ethanol, Sigma) was supplemented into the media upon isolation. Proliferating MuSCs were labeled by supplementing the media with 5- ethynyl-2’-deoxyuridine (EdU, 10 μM, ThermoFisher) for 2 hours prior to collection. For immunocytochemistry, myofibers were fixed with 4% PFA for 15 minutes at room temperature and then washed three times with PBS (Sigma) for 10 min each.

### Muscle stem cell isolation and culture

Hind limb muscles were dissected out of the mouse, mechanically diced, and enzymatically digested in collagenase (400 U/mL Worthington) in Ham’s F-12C (Gibco) for 60 min at 37°C with vigorous shaking every 10 min. Collagenase was quenched with 15% horse serum (Gibco) in Ham’s F-12C. The digest was passed through 100, 70, and 40 μm filters (Fisher Scientific) to remove debris and centrifuged (200 rcf for 5 min).

MuSCs were enriched by magnetic bead sorting using a Dead Cell removal kit (Milteyi) and satellite cell isolation kit (Miltenyi) following the manufacturer’s directions. The enriched MuSCs were resuspended in Ham’s F-12C (Gibco) media supplemented with Myocult^TM^ Expansion Supplement (Stem Cell Technologies) with penicillin-streptomycin (1% v/v, Gibco), and plated on plastic for 1 hour in 5% O_2_ and 5% CO_2_ at 37°C to remove adherent cells. After 1 hour, the enriched MuSCs were plated onto gelatin coated plastic and cultured in 5% O_2_ and 5% CO_2_ at 37°C for 1 passage. At ∼50% confluency, MuSCs were trypsinized for 2 min at 37°C and replated at 7,500 cells/cm^2^ on Matrigel coated coverslips and cultured until the designated time points. For myogenic differentiation, MuSCs were seeded at 25,000 cells/cm^2^ for low cell density and 125,000 cells/cm^2^ for high cell density. After 48 hours, the media was exchanged to low serum differentiation media (high glucose Dulbecco’s Modified Eagle’s Medium (DMEM) with horse serum (5% v/v, Life Technologies), penicillin-streptomycin (1% v/v, Gibco), and insulin-transferrin-selenium supplement (1x, ThermoFisher) and cultured for 4 days. For genetic YAP and TAZ knockout *ex vivo*, 4-hydroxytamoxifen (500 nM in ethanol, Sigma) was supplemented into the media for 24 hours. At time of collection for immunocytochemistry, the media was exchanged for 4% PFA for 15 minutes at room temperature and washed 3x in PBS for 10 min each.

### Immunocytochemistry

To fluorescently immunolabel proteins, fixed samples were permeabilized and blocked with 3% BSA (Sigma) and 0.25% Triton X-100 (Sigma) in PBS for 1 hour at room temperature. For proliferation assays, EdU was then conjugated to Alexa Fluor - 488 using the Click-iT EdU reaction kit (Molecular Probes) for 30 min. Primary antibodies in 3% BSA in 0.25% Triton X-100 were incubated overnight at 4°C. The following primary antibodies were used in these studies: mouse anti-Pax7 (1:750, Developmental Studies Hybridoma Bank), rabbit anti-MyoD (1:250, Santa Cruz Technologies), chicken anti-syndecan 4 (1:200, Cornelison et al., 2004), mouse anti-MF20 (1:250, eBioscience), rabbit anti-paxillin (1:250, abcam), and mouse anti-myogenin (1:250, F5D, Santa Cruz Biotechnology). Samples were washed 3x in PBS containing 0.05% Tween-20 for 10 min each and secondary antibodies, including Alexa Fluor -488, -555 and -647 (Invitrogen at 1:500), plus DAPI (1 μg/mL, Sigma) in 3% BSA and 0.25% Triton X-100 were incubated for 1 hour at room temperature. Rhodamine phalloidin (Cytoskeleton) was used at 1:1000 dilution. Samples were then washed 3x in PBS for 10 minutes each and left at 4°C until imaged. Myofiber samples were mounted in Mowiol supplemented with DABCO (Sigma) and stored at 4°C until imaged.

### TUNEL Assay

DNA fragmentation after knockout in MuSCs was assessed via the TMR red *in situ* cell death detection kit (Roche), following the manufacturer’s protocol. Briefly, fixed MuSCs were permeabilized with 0.1% Triton X-100 (Sigma) in 0.1% sodium citrate (Sigma) for 2 min on ice and then washed twice with PBS for 10 min each. For positive controls, DNase I (60 Kunitz, Qiagen) with 1% BSA (Sigma) in 50mM Tris-HCl (Sigma) was incubated for 10 min at room temperature. DNA breaks were then labeled using the TUNEL-reaction mixture for 1 hour at 37°C in a humidity chamber. Samples were washed three times in PBS and then followed by blocking and immunostaining detailed above.

### Fluorescent Image Acquisition

Muscle sections were acquired on a Nikon Spinning Disk Confocal (CSUX-A1, Yokogawa) equipped with an EMCCD camera (Ultra 888, Andor) with a 20x objective (N.A. 0.75). Focal adhesion images were taken using a Nikon A1R confocal microscope equipped with a 60x objective (N.A. 1.2 water immersion). For cell morphology, images were taken using an Operetta high-content confocal microscope with a 20x objective (Perkin Elmer) and analyzed using Harmony software (Perkin Elmer). All other fluorescent images were collected on a Zeiss LSM710 scanning confocal microscope with a 20x objective (N.A. 1.0). Images were visualized and analyzed with ImageJ.

### Cell Migration Assay

For evaluation of migration, MuSCs were seeded at 2,600 cells/cm^2^ on Matrigel-coated glass bottom well plates and cultured for 24 hours. Cells were then treated with either 4-hydroxytamoxifen (500nM in ethanol, Sigma) or ethanol for 24 hours prior to imaging. Before live-cell tracking, MuSCs were incubated with fresh media containing 1x SPY555-DNA (Cytoskeleton, Inc) for 3 hours to label the cells and then tracked continuously for 24 hours on a Nikon ECLIPSE Ti2 inverted microscope equipped with an Okolab environmental chamber. Images were acquired every 15 minutes. MuSC migration was quantified using Imaris software.

### Statistical Analysis

All statistical analyses were performed in Prism (GraphPad) with a Student’s t-test, one-way ANOVA, or two-way ANOVA as documented in each figure. At least three different biological replicates were used per study and *p* <0.05 was considered significant.

## Acknowledgments

This work was supported by grants from the NIH (DE016523 and DK120921) to KSA. NIH (AR049446 and AR070630) to BBO, and NIH Medical Scientist Training Program Grant T32GM008497.

